# Caracterization of A^m^ –A genomes chromosomes in diploid wheat, polyploid wheat and triticales by marker cytogenetic

**DOI:** 10.1101/2021.08.16.456506

**Authors:** Hammouda-Bousbia Dounia, Benbelkacem Abdelkader

**Author notes:** Auteur correspondant.

## Abstract

The distribution and the **Caracterization** of constitutive heterochromatin in A-A^m^ genomes of diploid wheat (progenitor), polyploid wheat (hybrids) and triticales (primary and secondary) are analyzed and compared by C-bands. The Comparison of zones rich in highly repeated DNA sequences marked by C bands on the all chromosomes of A^m^ - A genomes revealed an important structural heterogeneity. Four chromosomes of *Triticum monococcum* (1A^m^-3A^m^-4A^m^-5A^m^) are almost similar to their homologues in wheat (*Triticum durum, Triticum aestivum*) and triticale, by the presence or absence of C bands. Contrary to the chromosomes 2A^m^ (rich in heterochromatin),6A^m^-7A^m^ (absence of C bands) show a great differentiation compared to their homologues of *Triticum durum* and *Triticum aestivum* and x-*Triticosecale* Wittmack. In the triticales, A genome chromosomes are richer in heterochromatin compared to theirs homologous of polyploid wheats. This is explained by a “genome shock The confrontation of a C-bands genome (*Triticum monococcum*) with a C+ bands genome (durum wheat / or common wheat) produces an interspecific hybrid which at the sixth generation reveals C+ bands (triticales).The variations observed in our vegetal material indicated the existence of an intervarietal and interspecific heterochromatic polymorphism. The presence of B chromosomes in triticales, could be explained as a manifestation of their adaptation.

## Introduction

The genetic variability can be sought in wild species that constitute an important gene pool usable in the improvement of cultivated wheat such as *Titiccum. Dicoccum, Triticum urartu (*Liu et al., 2018) and *Triticum monococcum* L. (Megyeri et al., 2014). Einkorn wheat (*Triticum*.*monococcum* L.) with a unique genome constitution, 2n = 2x = 14, A^m^A^m^) (Megyeri et al.,2016) a food base for early farmers for several thousand years (Hidalgo and Brandolin.,2014) is one of the most ancient crops and the first domesticated wheat, It has been supposed that T. *monococcum* was domesticated from its wild progenitor T. boeoticum Boiss in southeastern Turkey, in the Karacadağ mountain range. (Heun et al., 1997) it was used in wheat breeding as a source of disease resistance genes, high carotenoid content, and other agronomically important characters (1997, Damania et al., 1998; Singh et al., 2008). Several tools for the analysis of genetic variability exist, some of which are based on morpho-physiological and cytogenetic criteria. The cytogenetic markers remain one of the essential analysis in the program of the wheat and the triticales selections, in order to highlight the anomalies which can affect the chromosomes during the crossings (Gill et al., 1991, Babaeva et al. 2007, 2015, Hammouda and Khalfallah, 2014, Hammouda et al 2017, 2021). The comparison of the genomes of several wheat species, from their wild forms to the cultivated forms used today, sheds light on original genetic mechanisms linked to the evolution of this cereal. (Li LF et al,, 2019).

The present work is part of genetic resources of the wheat and the triticales. We were interested in a comparative structural analysis of all A genomes, belonging to the : (i) the ancestral species *Triticum monococcum* (2n=2x=14, A^m^A^m^) et (ii) the polyploid wheats *(Triticum durum* (2n=4x=28, AABB, *Triticum aestivum* (2n=6x=42, AABBDD), iii) the primary triticale (2n=8x=56, AABBDDRR) and the secondary triticale (2n=6x=42, AABBRR) by a cytogenetic marker « C-bands **»**. In this study, the objective is to highlight :

‐ Identification and comparison of the all A^m^ chromosomes genome (progenitor) and A genomes in the studied species.
‐ Distribution and characterization of heterochromatic zones corresponding to highly repeated non-coding ADN sequences, by investigating the A^m^ genome introgressed into A genomes of polyploid wheats and triticales..
‐ Determination of the rôle of heterochromatin and B chromosomes in adaptation of studied species to unfavorable climatic conditions.

## MATERIALS AND METHODS

### 2.1 Materials

The plant material in the form of seeds is about four species:

‐ *Triticum monococcum* L. (progenitor, Genome A donor)
‐ *Triticum durum* Desf. (hybrid1, AABB formula genomic).
‐ *Triticum aestivum* L (hybrid2, AABBDD formula genomic).
‐ x*Triticosecale* Wittmack(Generation 1, AABBRR formula genomic)
‐ x*Triticosecale* Wittmack (Generation 2, AABBDDRR formula genomic).

List of species and studied varieties with their pedigree and origins are presented in the Table 1.

**Table 1.** The List and origins of the studied species.

| Species | Varieties / Lines | code | Ploidy level | Country of origin | Source | Pedigree |
| --- | --- | --- | --- | --- | --- | --- |
| *Triticum monococcum* (progenitor) |  |  | 2x |  | I.T.G.C |  |
| *Triticum durum* (Hybrid 1) | -Oued-zenati<br>-Cirta<br>Boussalem | O-Z<br>CI<br>BO | 4x | Syrie<br>Locals<br>Locals | I.T.G.C | Heider/Marli /Heider-Cro ICD 414—1BLCTR-4AP KB214-0KB-0KB-1KB-0KB |
| *Triticum aestivum* (Hybrid 2) | Manhondémias<br>Ziad<br>Tessalah | Ma<br>Zi<br>Te | 6x | Spaine<br>France<br>Mexico | I.T.G.C | PLC/RUFF/GTA"S"RoletteCm179 04 Alondra"S"ERA SONGLU//Alondra"S Norin10/Brevor//P14 |
| x-*Triticosecale* Witmack (Generation1) | Chrea<br>Chelia<br>Foca | Chr<br>Cli<br>Foc | 6x | Cimmyt<br>Cimmyt<br>Cimmyt |  | CTSS95Y00296S-10M-0Y-0B-0Y-0B-1B-<br>CTSS95Y00296S-10M-0Y-0B-0Y-0B-1B-<br>CTSS95Y00296S-10M-0Y-0B-0Y-0B-1B- |
| x-*Triticosecale* Witmack (Generation2) | Mahondemias X RC9<br>KVZ-alb x RC9<br>KVZ-alb x Landrace | MahxR<br>KvzxR<br>KvzxL | 8x | Locals<br>Locals<br>Locals | I.A.B | Mah x RC9<br>KVZ x RC9<br>KVZ x Landrace |

**I.T.G.C:** Technical Institute of Field Constantine. Algeria
**I.A.B:** Batna Institute of Agronomy
**CIMMYT:** International Maize and Wheat Improvement Center.

#### 2- Method

The used cytogenetic « C-banding » technique is described by Gill et al. (1991, 2009) on common wheat, Pignone et al (1989) on durum wheat and Badaev et al. (1992) on triticales. In the experimental procedures, we have modified hydrolysis, ADN-renaturation and coloration steps, by varying the concentrations of the solutions so as to have an optimum band coloration (Hammouda and Khalfallah, 2008, 2015; Hammouda et al. 2017).

The seeds are set to germinate in petri dishes on filter paper at a room temperature. Root tips are treated with ice water for 27h, fixed in the solution 3 ethanol: 1acetic acid (3v/1v), and then squashed en 45% acetic acid. The cover slips are removed with liquid nitrogen (–196°C). The best preparations have undergone the following steps in Table 2:

**Table2:** The different steps of C-banding applied to the four studied species.

|  | *Triticum monococcum* | *Triticum durum* Desf. | *Triticum aestivum* L. | *x-Triticosecale* Wittmack |
| --- | --- | --- | --- | --- |
| **Delamellation** | The detachment of the slides is made with liquid nitrogen (-196°) |  |  |  |
| **Deshydration** | All the slides are dried all over night. |  |  |  |
| **Hydrolyse** | none | The slides are immersed into HCl0.2 M at 60°C, 5 min. | The slides are immersed into HCl0.2 M at 60°C, 2 min. | none |
| **Denaturation of DNA** In a in barium hydroxide solution (formula):50g/ L | 4mn at 20°C. | 12 minutes room temperature | 7 minutes at room temperature | 6mn at 45°C |
| **Rinsing** | Tap water 30mn until clarification. |  | Distilled water and tap water until fully clarification |  |
| **Renaturation of DNA:** The slides are immersed in a solution Fresh 2Xssc (0.3M NaCl and 0.03 M Na citrate) at pH 7. | replaced by a hydrolysis with 1'HCl 1N at 60 °C | For 1 hour at 60°C. |  | 15minutes at 60°C, then, during 1h30 min at 52°C |
| **Giemsa coloration** In a Buffer Solution Sorensen phosphate at pH 6.8 | 10% for 30mn. | 6 % for 45mn | 4 % for 30 mn | 5% for 40 mn |
| **Mounting** | Slides are left dry all night, then fixed definitively with a mounting liquid « Depex ». |  |  |  |

## Results and Discussion

### Results

We have been able to identify all genomes of *Triticum monococcum* L. (A^m^A^m^), *Triticum durum* (AABB), *Triticum aestivum* (AABBDD) and thoses of primary triticale (AABBDDRR) and secondary triticale (AABBRR) (Figure. 1, mitotic chromosomes) (Figure 1).

**Figure 1:**
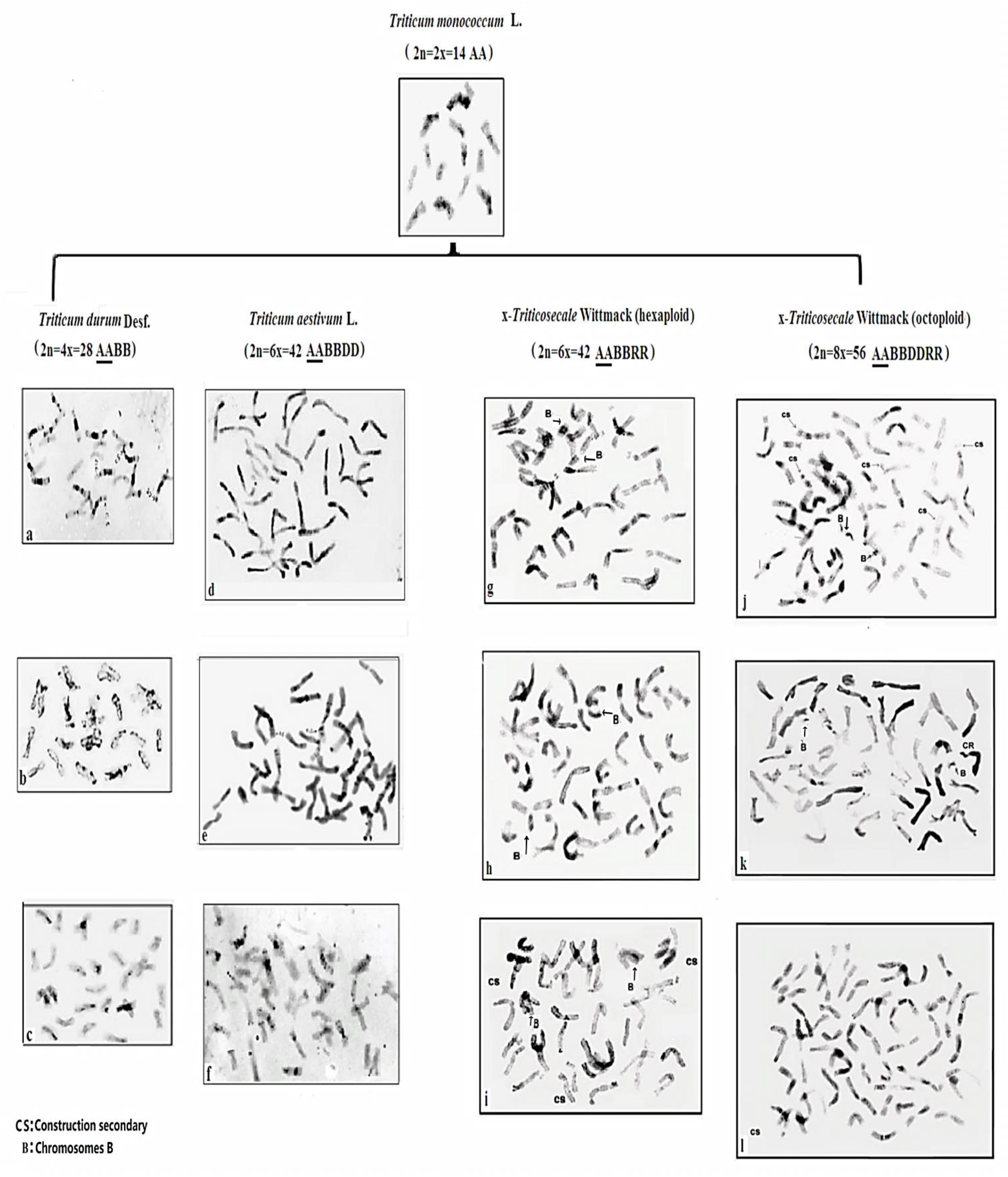
Mitotic chromosomes marked by C-banding: *Triticum monococcum, Triticum durum varieties* (Oued-Zenati(a), Cirta (b),Boussalem (c)),*Triticum aestivum* lines (Mahon-demias(d), Ziad(e),Tessalah(f)) and x-*Triticosecale* Wittmack variaties and lines (Chrea(g), Chelia(h), Foca(i), Mahon-demiasxRC9(j), KVZ/albxRC9(k), KVZ/ALBxLandrace(l)).

We remind that genome A donor is *Triticum monococcum* L. (Kuspira *et al*. 1986, ; Gill *et al*., 1987 ; Friebe *et al*. 1990. Ling et al., 2013.) which we are interested in studying its genome in comparison with their homologous of the studied species and varieties.

The identification and the distribution of constitutive heterochromatin (non-coding DNA sequences rich in CG bases) in the genomes (A - A^m^) of studied species are analyzed and compared by C-bands. This analysis revealed important differences in the zoning of chromosomes, in numbers and intensity of C+ additionals bands. Indeed, the number of bands and their localization on the chromosome, as well as their intensity, differs from one a species to another and one variety to another (Figure 1).

We remind that genome A donor is *Triticum monococcum* L. (Kuspira *et al*. 1986,; Gill *et al*., 1987 ; Friebe *et al*. 1990, Heslop-Harrison, 1992 ; Ling et al. 2013) which we are interested in studying its genome in comparison with their homologous of the studied species and varieties.

The identification and the distribution of constitutive heterochromatin (non-coding DNA sequences rich in CG bases) in the genomes (A - A^m^) of studied species are analyzed and compared by C-bands. This analysis revealed important differences in the zoning of chromosomes, in numbers and intensity of C+ additionals bands. Indeed, the number of bands and their localization on the chromosome, as well as their intensity, differs from one a species to another and one variety to another.

**In diploide wheat (**ancestor),

The analysis of C bands of genome A^m^ showed the fine bands marked on the majority of chromosomes, except chromosome 6A^m^. The 2A^m^ and 4A^m^ chromosomes revealed darck

Bands (figure 4).

**.Figure 3:**
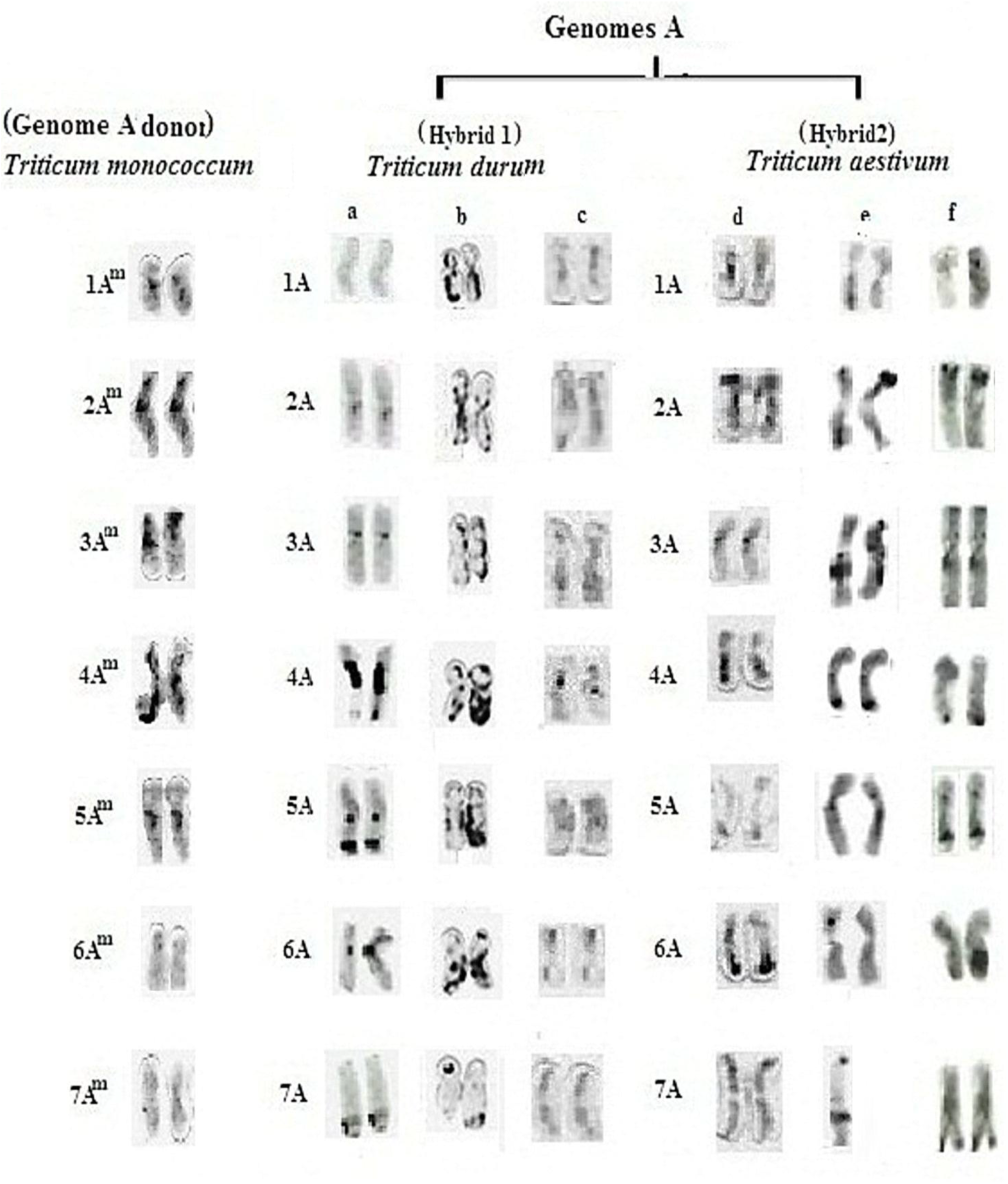
C-banding of chromosomes of A^m^ genome of diploide wheat, all A genomes of Durum wheat varieties (a-Oued-Zenati, b-Boussalem, c-Cirta) and commun wheat varieties (d-Mahondemias, e-Ziad, f-Tessalah).

**Figure 4:**
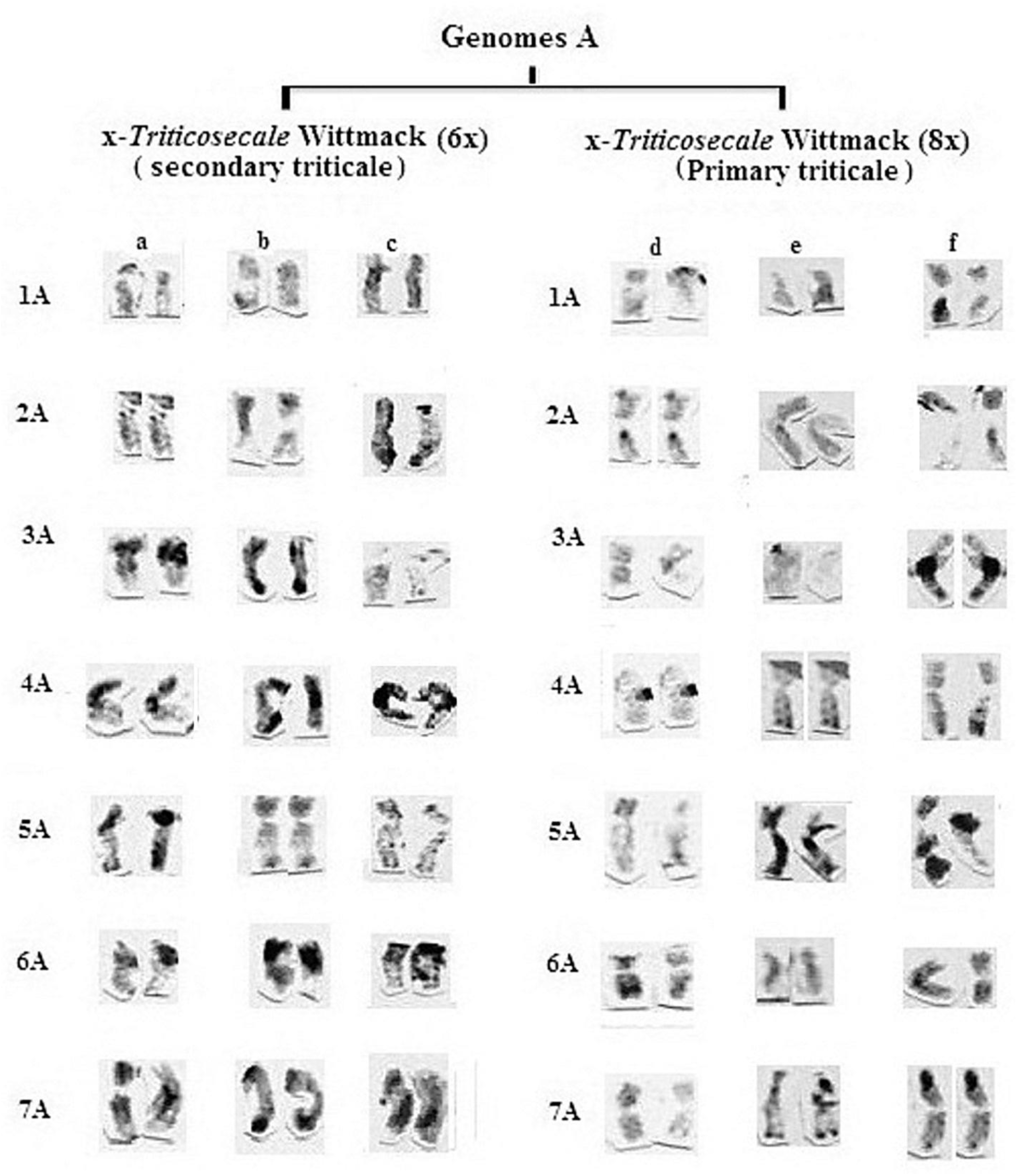
C-banding of chromosomes of A genomes of hexaploid triticale varieties (a-Chrea, b-Chelia, c-Foca) and octoploid triticale lines (d-MahondemiasxRC9, e-KVZ/albxRC9, f-KVZ/albxLandrace).

The Genomic formula:AA^m^ = 2x= 12 AA^m^ (metacentrics) + 2AA^m^ (Sub-metacentrics))= 14

The karyotipic formula: A = n= 6A^m^ (metacentrics) +1A^m^ (Sub-metacentrics)=7

#### In durum wheat. (Hybrid1)

In the Oued-Zenati variety, genomic analysis shows a overload of constitutive heterochromatin marked on 4A and 5A.chromosomes. The 2A 3A 6A 7A chromosomes revealed fines C bands, but, the 1A chromosome is unmarked (Figure 4, a)..

In the Cirta,variety, C-banding analysis of chromosomes showed variation in the additional bands distribution marked on the 1A, 2A, 4A, 5A, and 6A chromosomes(Figure 4, b)…

In the Boussalam variety, genomic analysis reveals the fines bands localized on 3A, 4A et 5A chromosomes. Howe ver, the 1A, 2A and 7A are unmarked. (Figure 4, c)..

#### In commun wheat. (Hybrid2)

The A genome of the Manhondemias variety is characterized by the presence of the additional dark bands (C+) marked on 1A, 2A, 3A, 4A, 6A chromosomes (Figure 3, d). its homologous genome of the Ziad variety reveals additional bands (C+) observed on chromosomes 2A, 3A, 4A, 5A and 6A Figure 3, e), and that of the Tessalah variety is poor in C+ bands, except 2A, 5A and 6A chromosomes (.(Figure3,f)

#### In hexaploid triticale (generation 1)

The A genome of the Chrea variety is displayed by the additional dark bands (C+) marked on all chromosomes except the 1A, 2A chromosomes (Figure 4, a). its homologous genome of the Chelia variety shows darck bands (C+) observed on the chromosomes 3A, 4A, 6A 7A, and fines bands on the 1A, 2A, 5A chromosomes Figure 4, b). That of the Foca variety is reveals C+ bands on the 2A, 4A, 6A and 7A chromosomes (.(Figure4,c).

#### In octoploid triticale (generation 2)

C-banding analysis of the A genomes of the Mah x RC9, KVZ x RC9, and KVZ x Landrace lines revealed many variations in chromosome zoning (figure 4, d,e,f). The additional darck bands (C+) are marked on 5A chromosome of KVZ/albxRC9, on 3A, 4A, 5A and 7A chromosomes of KVZ/albxLandrace (figure 4, e, f).

In addition, the B chromosomes of heterochromatic and / or euchromatic type are detected with in the triticales (Figure 1).

### Discussion

DNA denaturation-renaturation of C-banding denaturation and renaturation steps is critical because it needs an accurate pH, time and temperature for good heterochromatic band differentiation.

*Triticum monococcum* is an important source of useful genes and alleles that will be desirable for use in wheat selection programs. Well-defined Am chromosome markers would accelerate targeted introgression of T. *monococcum* chromatin in the wheat genome (Heslop-Harrison, 1992, Badaeva et al., 2015, ;Mégerie et al, 2014, 2017).

In this study, we recovered the distinct structural heterogeneity on the A^m^-A chromosomes. These heterogeneities are different between the sampled interspecifics hybrids (wheats and triticales) and their progenitor (*Triticum monococcum*

We only describe the variations of the additional C+ bands (absent in the A^m^ genome). We should note that for all (A^m^,-A) genomes, each type of chromosome is designated by a group:

#### 1A Group

The 1A^m^ chromosome is almost similar to their homologous wheat and triticale. Only, 1A chromosomes of the Cirta variety (durum wheat), the lines Mahondemias (common wheat) and Zvz/albxLandrace (octoploid triticale) showed the additionals bands (C+).

#### .2A Group

The 2A^m^ chromosome differs from their homologous of polyploid wheats and triticales by the presence of dark centromeric bands (long arm) and intercalary, which they do not have.

#### 3A Group

The 3A^m^ chromosome, compared to their 3A homologous of the varieties or lines studied (wheats and triticales), is distinguished by the presence or absence of C-bands. Only, 3A chromosomes of the varieties Boussalem (durum wheat) Chelia (hexaploid triticale) and Kvz/albxLandrace (octoploid triticale) which are characterized by additional C+ bands.

#### 4A Group

Only The 4A^m^ chromosome, which is characterized by heterochromatin overload (thick centromeric bands), which is observed on the 4A homologous of Oued-Zenati variety (durum wheat) and three varieties of the hexaploid triticale.

#### 5A Group

The 5Am chromosome, compared to their 5A homologous in wheat and triticales, is almost similar.

#### 6A and 7A Groups

The 6Am and 7Am chromosomes are different from their wheat and triticale 6A homologous 7A they are characterized by the absence of heterochromatic bands. Additionnal C+ bands are marked the majority of A chromosomes.

The Comparison of zones rich in highly repeated DNA sequences (Heterochromatin) marked by C bands on the all chromosomes of A^m^ - A genomes revealed a significant structural heterogeneity (Table **3**).

**Table 3:** C-band numbers of Am-A genomes of diploid wheat, polyploid wheat and triticale.

| Species |  | *Triticum monococcum* | *Triticum durum* |  |  | *Triticum aestivum* |  |  | x- *Triticosecale* (6x) |  |  | x- *Triticosecale* (8x) |  |  |
| --- | --- | --- | --- | --- | --- | --- | --- | --- | --- | --- | --- | --- | --- | --- |
| Genome (A<sup>m</sup>, A) |  |  |  |  |  |  |  |  |  |  |  |  |  |  |
| Varieties / Lines |  |  | O-Z | Cl | BO | Ma | Zi | Te | Chr | Cli | Foc | MahxR | KvzxR | KvzxL |
| chromosomes | 1A | 1 | 0 | 3 | 1 | 3 | 2 | 1 | 2 | 1 | 2 | 2 | 1 | 2 |
|  | 2A | 3 | 1 | 3 | 2 | 3 | 3 | 1 | 3 | 2 | 4 | 3 | 2 | 3 |
|  | 3A | 3 | 1 | 2 | 2 | 2 | 3 | 3 | 3 | 3 | 2 | 3 | 2 | 1 |
|  | 4A | 5 | 5 | 4 | 1 | 3 | 4 | 1 | 3 | 2 | 5 | 1 | 2 | 4 |
|  | 5A | 2 | 2 | 3 | 1 | 1 | 4 | 2 | 4 | 3 | 5 | 2 | 5 | 3 |
|  | 6A | 0 | 2 | 4 | 2 | 1 | 2 | 2 | 3 | 3 | 4 | 3 | 1 | 3 |
|  | 7A | 0 | 1 | 2 | 1 | 1 | 2 | 2 | 4 | 3 | 4 | 3 | 3 | 2 |
|  | Total of C-bands | 14 | 12 | 20 | 10 | 14 | 20 | 12 | 22 | 17 | 26 | 17 | 16 | 18 |
|  |  |  | 42 |  |  | 46 |  |  | 65 |  |  | 51 |  |  |

The obtained results demonstrate that the three chromosomes of *Triticum monococcum* (donor of the A genome), 2A^m^ (rich in C+ bands), 6A^m^ and 7A^m^ (no C bands), are squarely different from their 2A - 6A -7A homologous chromosomes in the varieties and the lines (wheat and triticale) (Figure 5, table 3).

**Figure 5:**
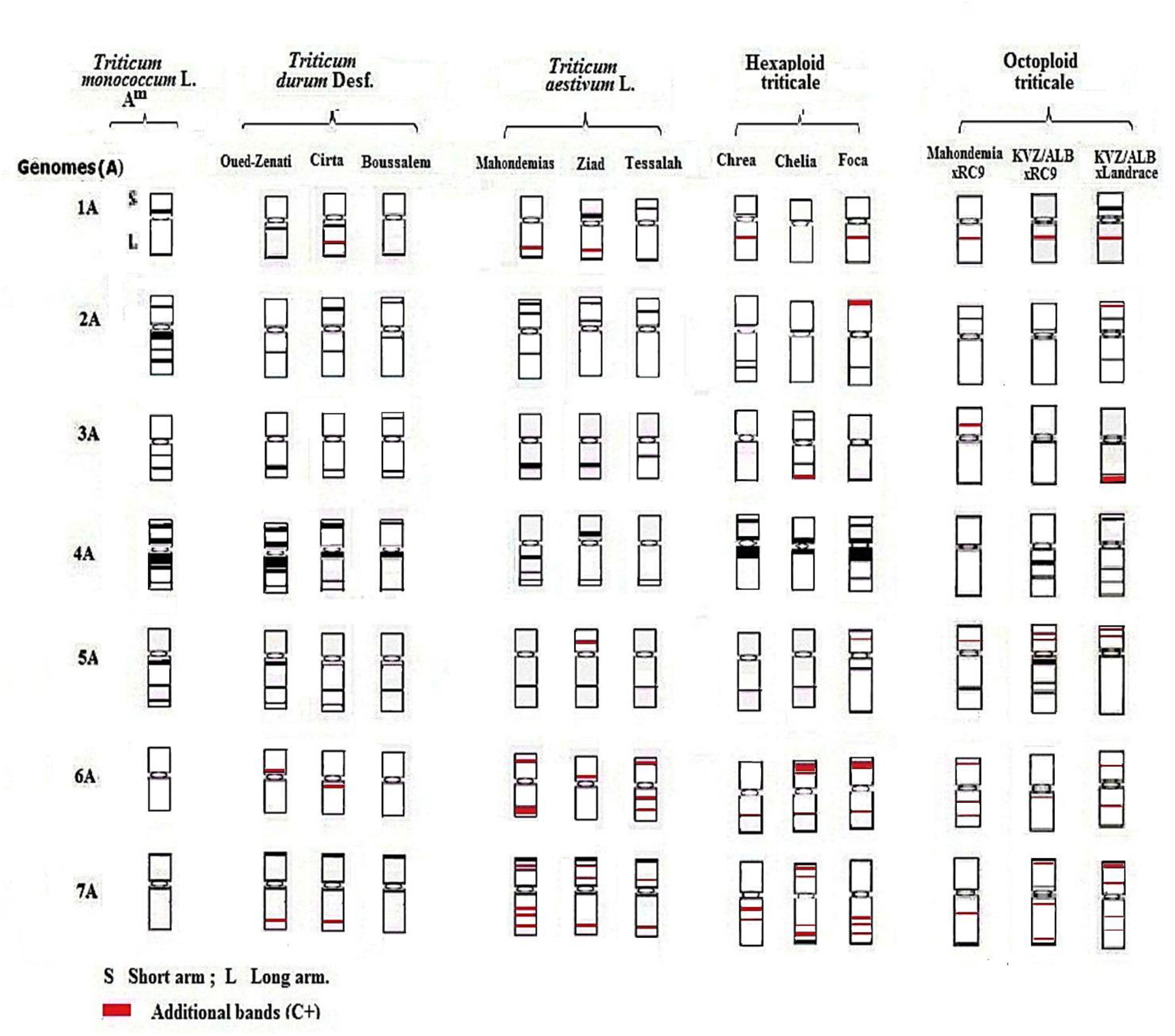
C-banded idiograms of A^m^ and A genome chromosomes. The most representative C-bands from each variety or line were used to construct the idiograms. 2A^m^ (rich in C+ bands), The 6A and 7A chromosomes of *Triticum monococcum* are unmarked (C-bands).

In the triticales, the chromosomes of A genome are richer in heterochromatin compared to theirs homologous A genomes of polyploid wheats. This is explained by a “genome shock » The confrontation of a C-bands genome (*Triticum monococcum*) with a C+ bands genome (durum wheat / or common wheat) produces an interspecific hybrid which at the sixth generation reveals C+ bands (triticales). The explicative hypothesis would be that the A genome of the triticales is enriched in heterochromatin during their stabilization. Figure 5, table 3).

All the variations revealed by the C-banding technique are probably due to the amplification or probably due to the amplification or reduction of the amount of highly repeated DNA sequences (rich in CG bases) in these regions.

Our obtained results in *Triticum monococcum*, polyploid wheats and triticales (secondary hexaploid, primary octoploid) in confrontation with those of authors (Badaeva et al., 2015 ;Mégerie et al (2014, 2017) working on *Triticum monococcum*, and common wheat, showing structural variations (position and intensity of C+ bands on the chromosome zoning).

Mégerie et al (2017), applied the technique of fluorescent in situ hybridization (FIH) couplet to microsatellites, on the chromosomes of *Triticum monococcum* – common t wheat showed different structural forms: the chromosomes 3Am and 6Am are similar with those of common wheat, The 2Am and 7Am chromosomes and squarely different. While, 1Am, 4Am and 5Am chromosomes are identical to their homologous common wheat, only chromosomes 2Am and 6Am are different from their 6A homologous wheat. Acording to The Work realized by Badaeva et al. 2015 on the chromosomes of diploid wheats (*T. monococcum, T. urartu* and *T. boeticum*), with the aim of determining the evolution of the chromosomes in the genus Triticum. In fact, they analyzed and compared the genomes A to their homologues of common t wheat,. They observed and detected NORs localized on chromosomes 1Au and 5Au Triticum Urartu, which are absent in *Triticum monococcum* and T*riticum boeticum*.

The same authors proved that Aesp_SAT86 (probe) can be used in the analysis of *Triticum monococcum* for discrimination and identification of particular chromosome species. According to Muégerie et al (2012), Microsatellite repeats facilitate introgression of Triticum monococcum chromatin into the polyploid background of wheat. Zhang et al. (2021) working on common wheat and its progenitors (*T. urartu*,

*A. speltoides*, and *Ae. tauschii*) have shown a series of genome rearrangements and sequence recombinations that have occurred, mainly in the heterochromatic regions of the chromosomes, which are an important marker for tracing genomic DNA sequence variations.

Other previous works (Sears., 1954), allowed the identification of all the A genomes in Tetraploid wheat and triticale by comparison made on the total lengths and ratios (BL/BC) with their homologues of the Chinese Spring variety. Gill et al (1991, 2009), Friebe and Gill (1994) propose, in the reference karyotype (ChineseSpring), to reverse the position of chromosomes 4A and 4B. Initially, chromosome 4B (old designation) does not appear to be a chromosome of the A genome due to unpairing with any chromosome of Triticum monococcum (Badaeva et al., 2007). According to Friebe and Gill (1994), the heterochromatin of chromosome 4B appears unstable. A pericentric inversion is detected in this chromosome in Chinese Spring (Endo et al, 2008, Wanda *et al* 1996; Silcova *et al*. 2007;Gill,2009,Kang *et al*, 2011; Méguéri et al 2012, Badaeva et al. 2015, Hammouda et *al*,, 2021).

In view of these considerations, the participants of the 7th I.W.G.S. voted on the distribution of chromosomes 4A and 4B. Thus, the former 4A is in genome B, designated as chromosome 4B. The former 4B is in genome A and designated as chromosome 4A. The constitutive heterochromatin, while non-coding, plays an important role in the regulation of the expression of the entire genome. Rearrangements in non-coding sequences, supposed to contribute to the genomic stabilization process, have been observed in synthetic polyploids of Triticeae (Ozkan et al, 2001; Shaked et al, 2001).

According to the authors (Stebbins 1971, Amirouche, 2007 ; Houben et al. 2011, Hammouda and Khalfallah, 2015, Hammouda et al.,2017, 2021), the apparition of B chromosomes is a form of adaptation of the species in difficult environment conditions. Our triticales have been cultivated in arid climate conditions. In this case, the presence of chromosomes B, could be explained as a manifestation of their adaptation.

## Conclusion

The cytogenetic markers remain one of the essential analysis in the program of wheat selections, in order to highlight the anomalies which can affect the chromosomes during the crossings. In this study we have tried to highlight the different chromosomal structural forms, by the intergenomic analysis (Am-A) of the four species Triticum monococcum L, *Triticum durum* Desf., *Triticum aestivum* L. and x-*Triticosecale* Wittmack (6x and 8x).

‐ The chromosomes of Triticum durum (F1 hybrid) and Triticum aestivum (F2 hybrid) and x-Triticosecale Wittmack (6x and 8x) are very rich in constitutive heterochromatin (specific bands) compared to their ancestors (*Triticum monococcum*).
‐ Four chromosomes of *Triticum monococcum* (1Am-3Am-4Am-5Am) are almost similar to their homologues in wheat (*Triticum durum* Desf, *Triticum aestivum* L) and triticale, and are characterized by the presence or absence of C bands. Contrary to the chromosomes 2A^m^ (rich in heterochromatin), 6A^m^ - 7A^m^ (absence of C bands) show a great differentiation compared to their homologues of durum wheat and common wheat and triticales.
‐ In the triticales, the chromosomes of A genome are richer in heterochromatin compared to theirs homologous A genomes of polyploid wheats in conclusion, *Triticum monococcum* L, constitutes an important reservoir of genes that can be used in the improvement of cultivated forms. The variations observed in the varieties and species studied indicated the existence of an intervarietal and interspecific heterochromatic polymorphism.

